# A High-Efficiency Geranylation Platform Enabled by Rational Tuning of a Single-Residue-Coded Chemo- and Regioselective Switch in Aromatic Prenyltransferases

**DOI:** 10.64898/2026.11.09.764520

**Authors:** Ruiying Qiu, Hong Zhang, Min Li, Zhihui Hu, Ziyi He, Shuo Ding, Longwei Gao, Yu Liu, Bo Chen, Man Huang, Zhi-Min Zhang, Jian-bo Wang

**Affiliations:** Key Laboratory of Multiple Organ Failure (Zhejiang University), Ministry of Education, Department of General Intensive Care Unit of the Second Affiliated Hospital, Zhejiang University School of Medicine, Hangzhou, 310058, P. R. China; Department of Microbiology and Institute of Pharmaceutical Biotechnology, Zhejiang University School of Medicine, Hangzhou, 310058, P. R. China; Key Laboratory of Chemical Biology and Traditional Chinese Medicine Research (Ministry of Education) and Key Laboratory of Phytochemistry R&D of Hunan Province, College of Chemistry and Chemical Engineering, Hunan Normal University, Changsha, 410081, P. R. China; State Key Laboratory for Quality Ensurance and Sustainable Use of Dao-di Herbs, Beijing, 100700, P. R. China; State Key Laboratory of Bioactive Molecules and Druggability Assessment, Jinan University, Guangzhou, 511436, P. R. China; Key Laboratory of Tea Science of Ministry of Education, College of Horticulture, Hunan Agricultural University, Changsha, 410128, P. R. China; Collaborative Innovation Center of Yangtze River Delta Region Green Pharmaceuticals, Zhejiang University of Technology, Hangzhou 310014, P. R. China; Binjiang Institute of Zhejiang University, Hangzhou, 310053, P. R. China; College of Life Sciences, Zhejiang University, Hangzhou, 310058, P. R. China

**Keywords:** Prenyltransferase, Geranylation, Directed evolution, Chemoselectivity, Regioselectivity

## Abstract

Geranylation is an efficient post-modification strategy for enhancing or introducing new properties to molecules; however, achieving both high efficiency and precision remains challenging. Aromatic prenyltransferases (PTs) provide a mild and sustainable approach, yet their practical application is often limited by the trade-off between broad substrate scope and low selectivity, and successfully balancing these properties is rare. Here, using the pharmaceutically relevant cannabigerol framework as a model, we identify a single-residue switch in a prenyltransferase from *Aspergillus terreus* (AtaPT) that governs both chemo- and regioselectivity. Rational tuning of this switch preserves broad substrate scope while enhancing selectivity, enabling the selective formation of each of the three possible mono-geranylated products in the model reaction. Structural analysis reveals that this residue modulates the selectivity and activity through coupled steric and electrostatic effects. Notably, transplantation of this site into three homologous PTs similarly reprogrammed selectivity, establishing a conserved and generalizable control mechanism. Leveraging these insights, we achieved selective geranylation across 36 aromatic substrates, affording 41 products. Integration with an efficient geranyl donor regeneration system enabled preparative-scale synthesis, reaching titers up to 324 mg/L. This work defines a unifying mechanism for selectivity control in PTs and provides a high - efficiency platform for geranylation.

## 1 Introduction

Prenyltransferases (PTs) catalyze prenylation across a wide range of biomolecules—including proteins, nucleotides, and small organic molecules—serving as key biocatalysts for this important modification [1-3]. Owing to its ability to significantly enhance membrane permeability and bioactivity [4-6], prenylation is a highly valuable transformation. PTs offer a direct and mild route to achieve it, making them promising catalysts for diverse applications, especially in pharmaceutical sciences. Among various PTs, soluble aromatic prenyltransferases (aPTs) have attracted growing attention due to their exceptional catalytic performance [7, 8]. These enzymes accept diverse donors, ranging from natural C5–C20 substrates (e.g., DMAPP, GPP, FPP, GGPP) to non-natural analogs with varied carbon chains or phenyl substituents. Their acceptor scope is similarly broad, encompassing flavonoids, indoles, alkaloids, and peptides [9, 10]. However, most reported aPTs face a fundamental trade-off: high selectivity often correlates with narrow substrate specificity, whereas broad substrate scope tends to coincide with low selectivity. Balancing these catalytic properties is therefore crucial to advance the practical utility of aPTs.

Driven by the declining costs of gene synthesis and sequencing, numerous aPTs with broad substrate scope have been reported, providing a rich repository of enzymes to address the aforementioned balance between substrate promiscuity and high selectivity [11-18]. Elucidating the selectivity-determining mechanisms of PTs is crucial for designing robust variants that combine broad substrate-scope with high fidelity. However, studies that directly investigate the intrinsic selectivity regulation in aPTs remain scarce. To the best of our knowledge, mechanistic and engineering efforts for PTs have largely focused on the relatively well-characterized dimethylallyltryptophan synthase (DMATS) family [19, 20], which primarily acts on simple indole substrates [21-23]. Furthermore, current engineering strategies often rely on a template-driven approach, where modifications are inferred from comparative analysis of enzymes with divergent selectivity patterns [21, 24]. Notably, precise and rational control over both chemo- and regioselectivity in PTs has not yet been achieved. Moreover, previously proposed mechanisms for regioselectivity are largely attributed to multisite synergistic effects, rather than a well-defined single-residue determinant (Figure 1).

**FIGURE 1.**
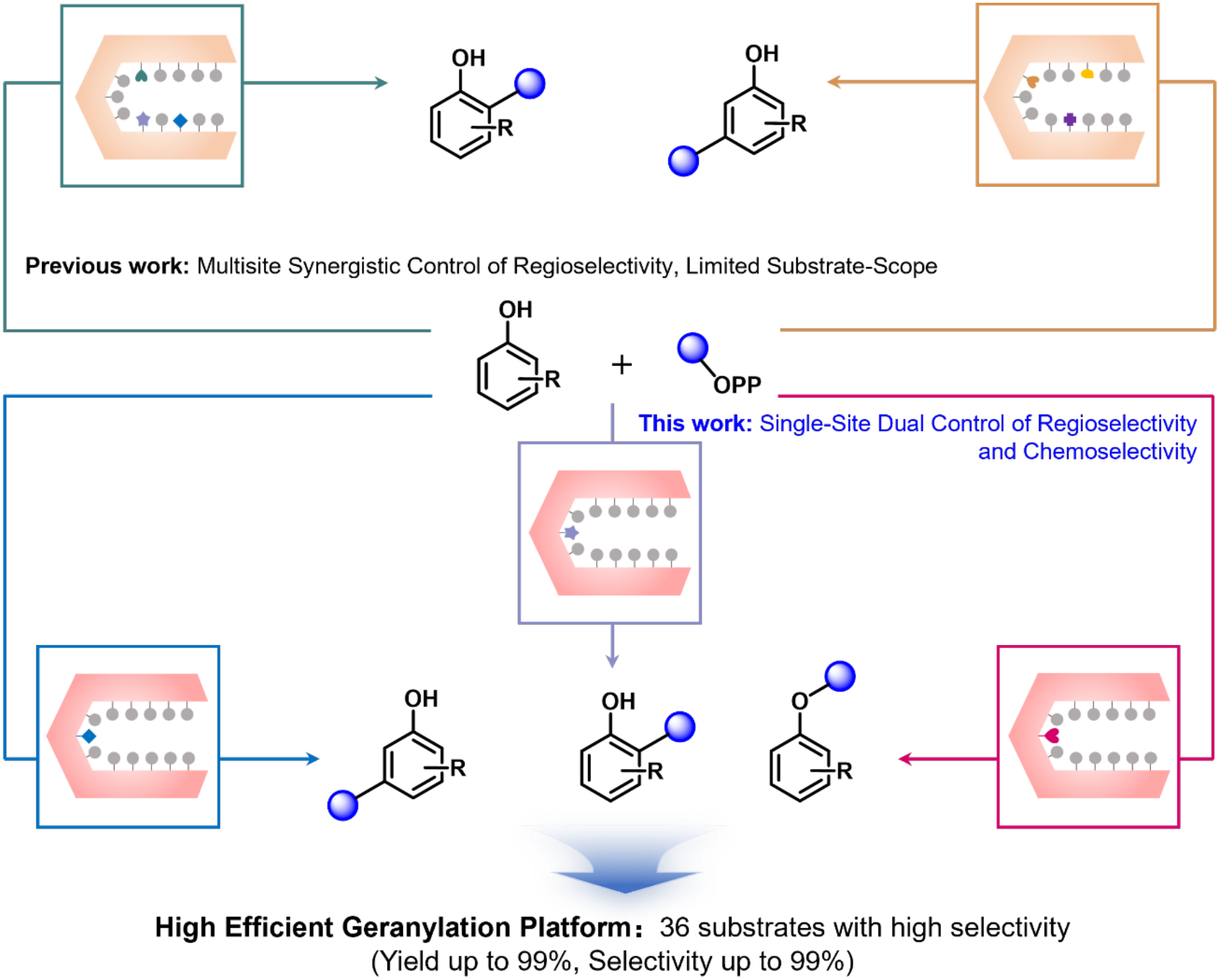
Mechanistic investigation of regio- and chemoselectivity in aromatic prenyltransferases.

Here we selected the aromatic prenyltransferase AtaPT [11] from *Aspergillus terreus* as a model aPT, owing to its well-documented promiscuity toward diverse prenyl donors (C5, C10, C15, C20) and acceptors (including lignans, tryptophan dipeptides, quinoline alkaloids, coumarins, flavones, and flavonoids), along with its ability to generate varied selectivity outcomes—such as single- and multi-site *O*- and *C*-prenylations in multiple substitution patterns. Elucidating the mechanism governing selectivity toward defined substrate types would provide a strategy to balance broad substrate scope with high selectivity. To this end, we chose olivetol and geranyl pyrophosphate (GPP) as model substrates. Using directed evolution [25-30], we successfully identified a single residue that efficiently controls both regio- and chemoselectivity. Further, through structural resolution, we decoded the mechanistic basis underlying activity and selectivity regulation. Finally, by expanding the substrate scope and achieving the synthesis of 13 geranylated polyphenols via an *in situ* GPP regeneration system, we established a general and scalable biocatalytic platform for the selective geranylation of small-molecule polyphenols.

## 2 Results and Discussion

### 2.1 Testing the Activity of the AtaPT-WT toward the Model Reaction

Olivetol (**1a**) can theoretically undergo monogeranylation at three distinct positions, yielding an *O*-geranylated and two *C*-geranylated products. To assess the inherent selectivity of AtaPT toward this substrate, whole cells expressing AtaPT-WT were employed as catalysts. Reaction mixtures were analyzed by HPLC and LC-MS.

As shown in Figure 2a and Figure S1, all three possible monogeranylated products were formed with a total yield > 40%, confirming the promiscuous selectivity of AtaPT-WT. Among these, the *C*2-geranylated derivative, commonly known as cannabigerol (CBG, **2a**), exhibits a broad range of reported bioactivities including antibacterial, antifungal, anticancer, anti-inflammatory, redox-modulating, and neuromodulatory effects [31-34]. Comparison with a commercial CBG standard revealed that **2a** was produced with low efficiency: yield was below 10%, and it accounted for only 20.6% of the total product distribution (Figures 2a and S1).

**FIGURE 2.**
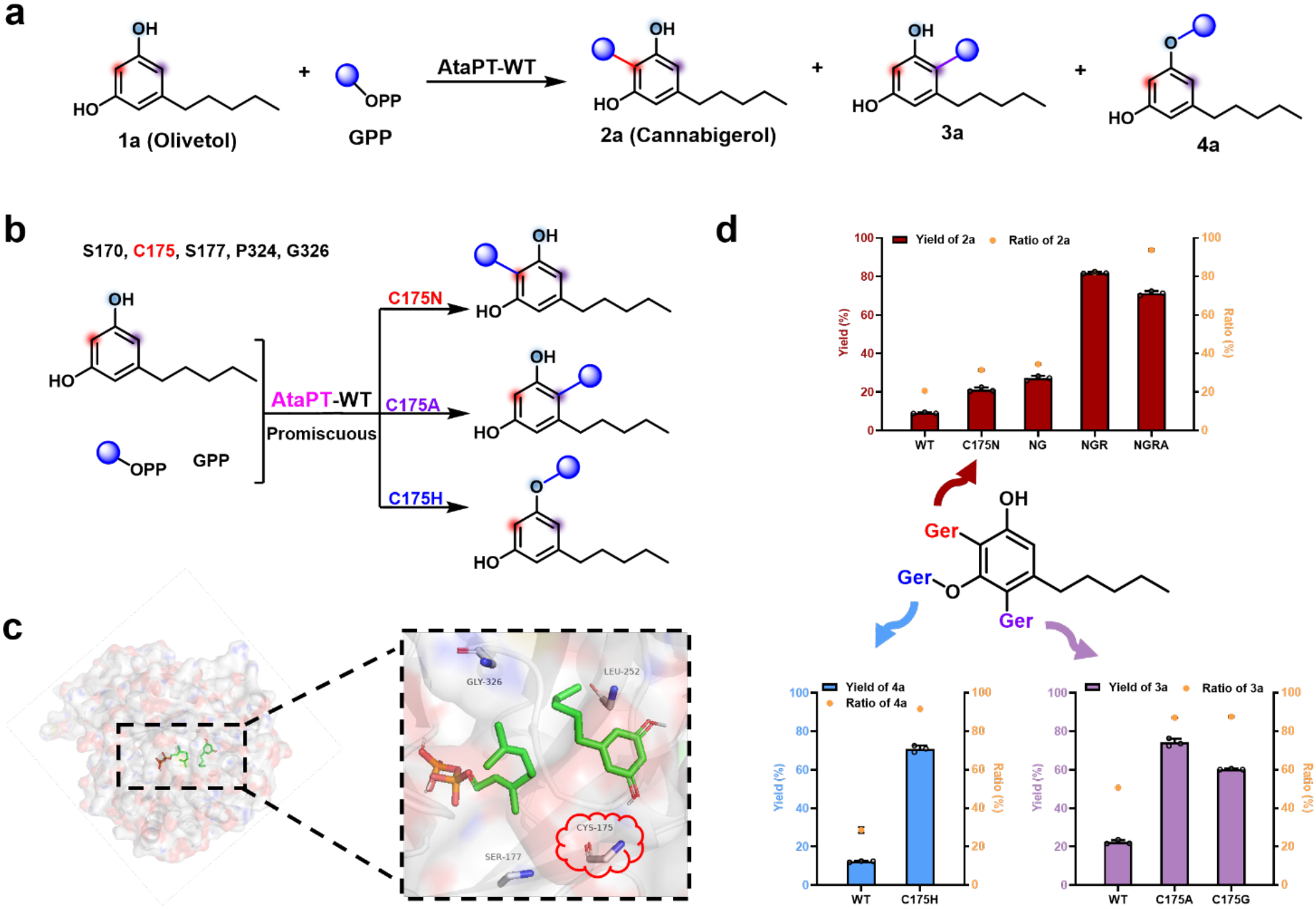
Evolution of AtaPT for efficient synthesis of geranylated olivetol derivatives. (a) Geranylation of **1a** catalyzed by AtaPT-WT using GPP as the donor. (b) Site-saturation mutagenesis (SSM) of AtaPT toward model substrates revealed that C175 is a key residue. (c) Key mutated residues in substrate binding pocket. Substrate **1a** and GPP are shown as green sticks, and key sites (C175, S177, L252, and G326) are shown as gray stick. (d) Geranylation of **1a** catalyzed by the recombinant AtaPT mutants using GPP as donor. Improvement in catalytic activity across four rounds of directed evolution using SSM and iterative saturation mutagenesis (ISM) for **2a**. Geranylation activity of AtaPT variants toward the model substrate **1a** for the production of **3a** and **4a**.

The other two products were assigned based on polarity considerations. The *O*-geranylated isomer, expected to be less polar, displayed a longer retention time in reversed-phase chromatography, allowing its identification as **4a**. Consequently, the remaining product **3a** was assigned as the *C*4-geranylated olivetol derivative (Figures 2a and S1).

While **2a** and **3a** occur only in trace amounts in *Cannabis sativa*, efficient and highly selective synthetic methods for them remain unavailable, and **4a** has not yet been characterized. Engineering AtaPT variants with high selectivity toward each of these products would therefore not only provide mechanistic insights into enzymatic regiocontrol, but also enable the systematic study and application of these previously inaccessible geranylated phenols.

### 2.2 Directed Evolution of AtaPT

Building on prior mutagenesis studies [35], we first performed site-saturation mutagenesis (SSM) [36] at five key residues—S170, C175, S177, P324, and G326—previously implicated in modulating activity and selectivity. Mutants were screened using whole-cell biotransformation in 96-deep-well plates, with products analyzed by HPLC. While libraries at P324 and G326 yielded no improved variants, several mutations at S170, C175, and S177 enhanced performance relative to the wild type.

For the target product **2a**, improved activity and selectivity were observed for mutants S170P (yield: 13.7%; selectivity: 19.1%), S170T (yield: 11.7%; selectivity: 17.3%), C175L (yield: 10.7%; selectivity: 26.4%), C175I (yield: 9.9%; selectivity: 24.6%), S177M (yield: 20.7%; selectivity: 28.0%), and S177Q (yield: 16.3%; selectivity: 19.9%) (Figure S2). For the 4 - *C*-geranylated product **3a**, C175A (yield: 74.1%; selectivity: 87.1%) exhibited superior activity and selectivity. For the *O*-geranylated product **4a**, enhancements were achieved with C175H (yield: 70.9%; selectivity: 91.6%).

The pronounced effect of mutations at C175 identified this residue as a key switch governing both regioselectivity and chemoselectivity. We therefore constructed and analyzed all 20 variants at this position (Figure S3). Clear selectivity trends emerged: small residues (e.g., Ala, Gly) strongly favored **3a**, with C175A proving optimal. Medium-sized residues (e.g., His, Leu, Gln) shifted preference toward **4a**, with C175H as the top performer. Given their excellent catalytic performance toward **3a** and **4a**, respectively, C175A and C175H were selected as the final variants for subsequent mechanistic studies and biomanufacturing. For the desired **2a**, the best single variant was C175N, which provided 21.2% yield and 31.4% selectivity, indicating clear potential for further optimization (Figure 2b).

To further improve the catalytic performance toward **2a**, we subjected the C175N variant to additional rounds of directed evolution. Since residues S170 and S177 both appeared beneficial for activity and selectivity—with mutations at S177 conferring more pronounced enhancements—we first performed iterative saturation mutagenesis (ISM) [37-39] at position S177. This round yielded the double mutant C175N/S177G (NG), which showed only modest improvements in both activity and selectivity (yield: 27.2%; selectivity: 34.5%). Using NG as the template, we then introduced targeted mutations at S170, but no variant with further improved performance was obtained. Since the selectivity of the NG mutant toward **2a** remained unsatisfactory, an additional round of ISM was warranted. To identify further mutation hotspots, we sequentially docked **1a** and GPP into the crystal structure of wild-type AtaPT (PDB: 5KCG) [11]. Based on the resulting model (Figure 2c), iterative mutations were introduced at two positions using NG as the template: N172 (located within the same loop as S170, C175, and S177) and L252 (positioned near **1a** on the same face as C175). While mutant N172H/C175N/S177G showed only marginal improvement in selectivity with unchanged activity (44.0%), mutant C175N/S177G/L252R (NGR) delivered a substantial increase in both activity (81.9%) and selectivity (82.4%). To further increase selectivity, we performed ISM on three additional residues using NGR as template: G251 (adjacent to **1a** and L252), A270 (opposite C175 and near **1a**), and G326 (opposite C175 and close to GPP). Although the corresponding quadruple mutant showed higher selectivity at the cost of activity, the quadruple mutant C175N/S177G/L252R/G326A (NGRA) achieved an optimal balance, exhibiting high activity (71.3%) and excellent selectivity (93.7%) toward **2a** (Figure 2d).

To determine whether more efficient evolutionary pathways could yield the final mutants and to identify key mutations governing activity and selectivity, we performed complete deconvolution [40] analysis (Figure S5). The results confirm that the pathway we employed indeed represents the most efficient route to the final variant, with C175 acting as the primary hotspot that critically determines both activity and selectivity. S177 also contributes to enhanced catalytic performance, consistent with earlier observations. The introduction of G326 notably improved selectivity, albeit with some reduction in activity. Interestingly, while the L252R mutation alone completely abolished activity, its combination with C175N and S177G led to a significant boost in both activity and selectivity, suggesting strong synergistic interactions among L252, C175, and S177.

Following the successful generation of four functional mutants—NGR, NGRA, C175A, and C175H—each capable of catalyzing the formation of distinct *C*- or *O*-geranylated olivetol derivatives, reaction conditions were optimized (Figures S6–S10), followed by detailed structural characterization. The results indicated, as expected, that **2a** is a *C*2-geranylated Olivetol, identified as CBG; **3a** is geranylated at the *C*4 position, whereas **4a** is geranylated at the 3-*O* position (Figure S11).

### 2.3 Kinetic Characterization of AtaPT Variants

To elucidate the catalytic basis underlying the enhanced performance of the NGR, NGRA, C175A, and C175H variants, steady-state kinetic assays were performed using **1a** as the acceptor substrate (Table S8 and Figures S12-14). Although the WT enzyme exhibited a relatively high turnover number (*k*_cat_ = 1.72 min⁻¹), it also showed a large *K*_m_ (1.59 mM), resulting in modest catalytic efficiency (*k*_cat_/*K*_m_ = 18.03 s⁻¹·M⁻¹). In contrast, the NGR variant displayed a lower *k*_cat_ (0.95 min⁻¹) but a markedly reduced *K*_m_ (0.47 mM), leading to improved catalytic efficiency (*k*_cat_/*K*_m_ = 33.69 s⁻¹·M⁻¹). Both NGRA and C175H showed slight decreases in *k*_cat_ and *K*_m_ relative to WT, yet their catalytic efficiencies (21.30 and 20.55 s⁻¹·M⁻¹, respectively) remained moderately enhanced. Notably, C175A combined an increased *k*_cat_ (1.94 min⁻¹) with a reduced *K*_m_ (0.90 mM), yielding a catalytic efficiency (35.93 s⁻¹·M⁻¹) nearly twofold higher than that of WT. These results indicate that the enhanced activity primarily arises from increased substrate affinity, with the C175A mutation improving both substrate binding and overall catalytic performance.

### 2.4 Mechanistic insights into the selectivity of AtaPT

To decipher the chemo- and regioselective mechanism of AtaPT toward the model reaction, we first generated crystals of all four variants in their apo form (Figures 3 and S16). These crystals were subsequently soaked with GSPP/GPP and **1a**. Ultimately, we determined four distinct complex structures: C175A‒GSPP‒**1a** (PDB: 27RS), C175H‒GSPP‒**1a** (PDB: 27SJ), NGR‒GPP‒**1a** (PDB: 27UF), and NGRA in complex with pyrophosphate (PDB: 27VQ), at resolutions of 2.5 Å, 2.1 Å, 2.9 Å, and 2.8 Å, respectively (Table S9). Of note, the long aliphatic tail of **1a** exhibits substantial structural flexibility. We therefore modelled olivetol primarily based on the well-defined electron density of its aromatic ring (Figure S16). Structural superimposition of these mutants revealed that C175 is located on the distal side of the geranyl-binding pocket within the *β*-barrel that accommodates both the substrate and the geranyl donor (Figure 3). No global conformational rearrangements were observed among the variants, suggesting that substitutions at C175 regulate AtaPT catalysis primarily by fine-tuning the local chemical environment.

**FIGURE 3.**
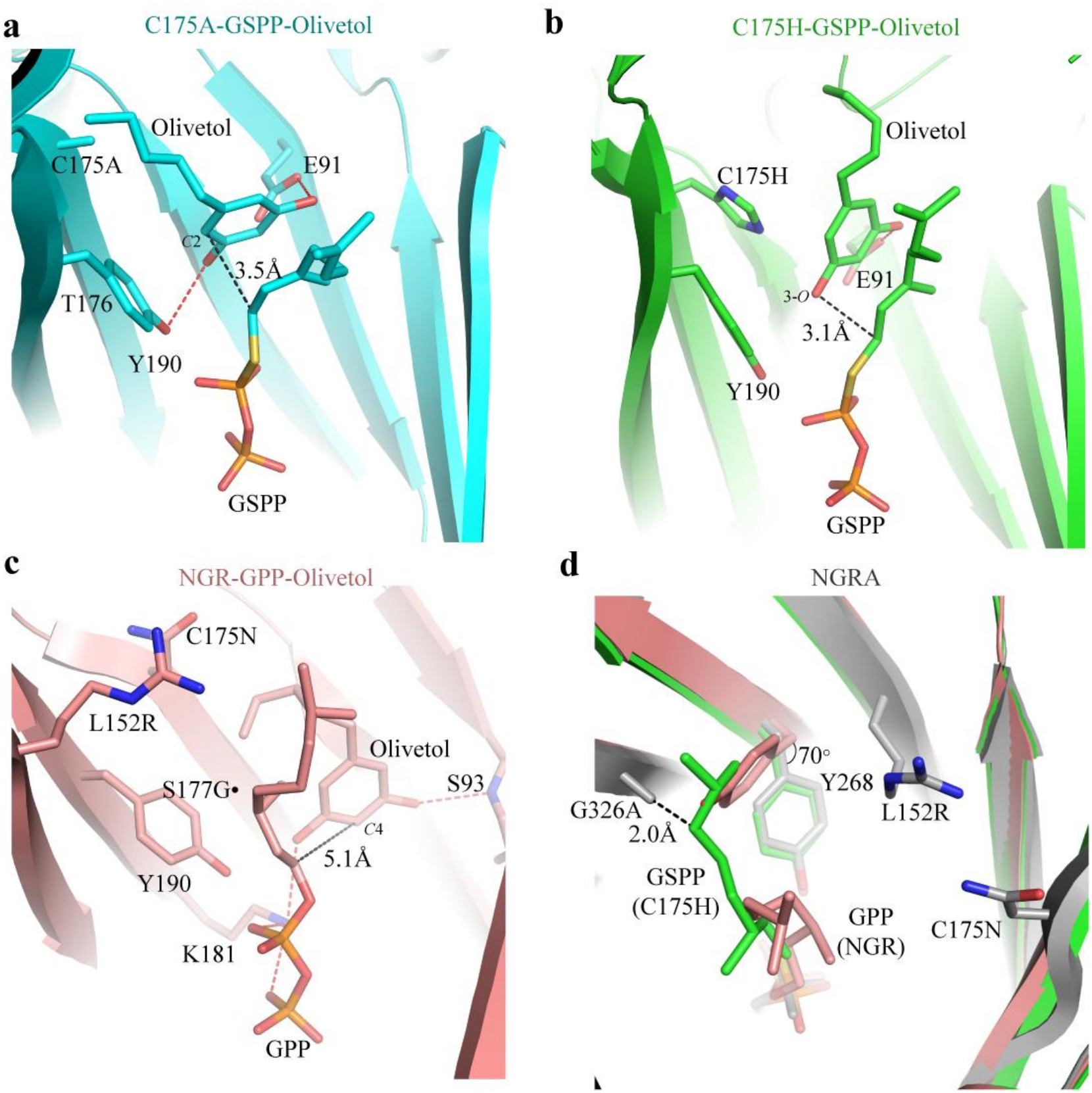
Crystal structures of AtaPT mutants. (a-c) Close-up views of the olivetol-binding pocket in C175A (a), C175H (b) and NGR (c). **1a**, GSPP/GPP, and adjacent residues are labeled and shown as sticks, with hydrogen bonds illustrated as red dashed lines. Distances from the 2-*C*, 3-*O* and 4-*C* of **1a** to the electrophilic carbon of GSPP/GPP are 3.5 Å, 3.1 Å and 5.1 Å, respectively. S177G mutation site in NGR was shown as black dot. (d) Superposition of the C175H, NGR and NGRA structures. The minimal distance between residue G326A and bound GSPP in the C175H mutant is 2.0 Å.

In the C175A mutant, the smaller alanine residue generates additional cavity volume to accommodate the aliphatic tail of substrate **1a**. Two hydrogen bonds are formed between the hydroxyl groups on the aromatic ring of **1a** and the side chains of Y190 and E91. Together, these interactions orient the *C*2 position of the benzene ring toward the electrophilic carbon of GSPP at a distance of 3.5 Å. This geometry strongly favors formation of the *C*2-geranylated product 3a (Figure 3a).

In contrast to C175A, the C175H mutant introduces a bulkier imidazole group adjacent to the substrate-binding pocket, which sterically shifts **1a** away from the mutation site (Figure 3b). A hydrogen bond is formed between the 1-hydroxyl group of **1a** and the side-chain of E91. These interactions poise the 3-hydroxyl group in close proximity to the electrophilic carbon atom of the geranyl moiety at a distance of 4.2 Å, accounting for the favored 3-*O* geranylation observed in the C175H mutant.

In the NGR variant, replacement of C175 with asparagine induces a conformation distinct from that observed in C175H, with the asparagine side chain flipping away from GPP (Figure 3c). Meanwhile, the S177G substitution removes side-chain bulk, collectively allowing substrate **1a** to access a deeper region of the *β*-barrel. In this configuration, the two hydroxyl groups of **1a** form hydrogen bonds with K181 and the backbone of S93, respectively. These interactions position the C4 site of **1a** toward the electrophilic carbon of GPP at a distance of ∼5.1 Å, consistent with the observed catalytic outcome. Notably, K181 also interacts with the negatively charged pyrophosphate moiety of GPP, suggesting that it may facilitate catalysis by acting as a general base.

Unlike the buried C175N and S177G mutations, L252R is located near the geranyl tail of GPP. The elongated arginine side chain induces an approximately 70° rotation of Y268, which in turn displaces the geranyl group from its canonical binding mode—similar to the conformations observed in the C175A and C175H mutants—and redirects it toward the center of the pocket, potentially constraining GPP for productive transfer (Figure 3d). Although a structure of the NGRA mutant in complex with GPP and **1a** was not obtained, the positioning of the G326A mutation directly on the geranyl-binding surface suggests a role analogous to that of the rotated Y268 in the NGR variant (Figure 3d).

In summary, structural analyses reveal that mutations at C175 precisely tune catalytic specificity by subtly remodeling both the substrate-binding pocket and the GPP-bound conformation.

### 2.5 Mechanism-guided design of PTs beyond AtaPT

The catalytic role of the key residue C175 has been elucidated; however, it remains important to determine whether its function can be transferred to other aPTs to guide the rational mining and design of enzymes with target selectivity. In our previous work [35], three homologous enzymes—AfaPT (94.81% identity), AaaPT (86.70%), and AmaPT (56.87%)—were identified, all of which contain the conserved XXLPXX motif and a highly similar loop region (residues 167–170) (Figure S17). Based on this, further rational engineering of these homologous enzymes was undertaken.

Initially, the wild-type activities of these enzymes were evaluated in the olivetol geranylation reaction (Figure 4a). Both AfaPT-WT and AaaPT-WT exhibited moderate activity (< 40%) and predominantly produced **3a**, with activities of 22.4% and 23.1% and selectivities of 56.0% and 54.6%, respectively. In contrast, no detectable product was observed for AmaPT-WT. The engineered variant AmaPT-SLG [35], in which the active pocket was remodeled, showed a notable improvement in activity, while **3a** remained the major product (activity 16.7%, selectivity 42.7%). Nevertheless, the overall activity and selectivity of AfaPT-WT, AaaPT-WT, and AmaPT-SLG remain suboptimal, underscoring the need for further engineering (Figure 4a).

**FIGURE 4.**
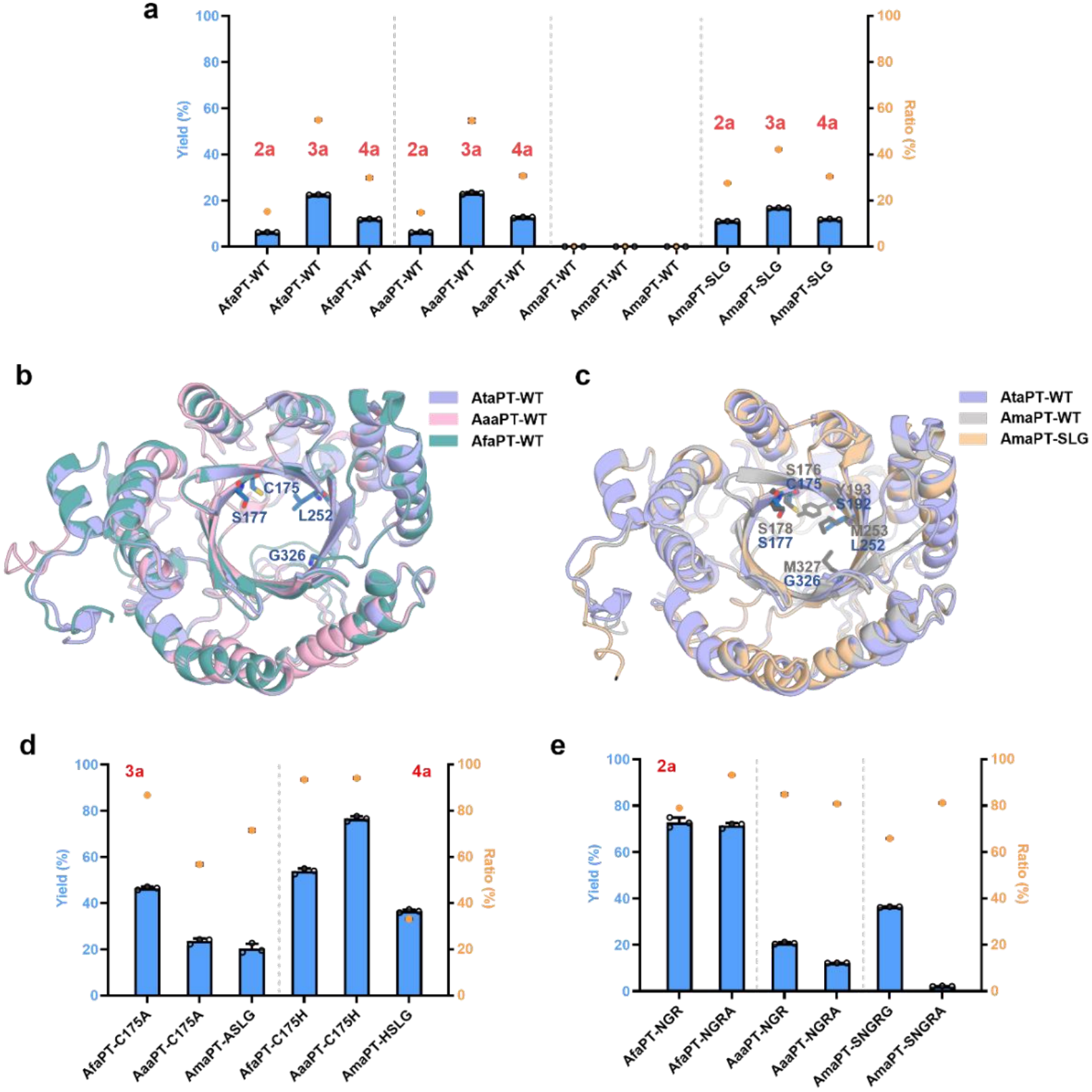
Mechanism-guided design of PTs beyond AtaPT. (a) Activity assays of AfaPT-WT, AaaPT-WT, and AmaPT-WT toward **1a**. (b) Structure alignment of AtaPT-WT with AfaPT-WT and AaaPT-WT. (c) Structure alignments of AtaPT-WT with AmaPT-WT and AmaPT-SLG. (d) Activity assays of AfaPT-C175A, AaaPT-C175A, AmaPT-ASLG, AfaPT-C175H, AaaPT-C175H and AmaPT-HSLG toward **1a** with GPP. (e) Activity assays of AfaPT-NGR, AfaPT-NGRA, AaaPT-NGR, AaaPT-NGRA, AmaPT-NGSRG and AmaPT-NGSRA toward **1a** with GPP. The crystal structure of AfaPT-WT, AaaPT-WT, AmaPT-WT and AmaPT-SLG was predicted by AlphaFold3. WT: wild type, AmaPT-SLG: Y193S/M253L/M327G, AmaPT-ASLG: Y193S/S176A/M253L/M327G, AmaPT-HSLG: Y193S/S176H/M253L/M327G, AfaPT-NGR: C175N/S177G/L252R, AfaPT-NGRA: C175N/S177G/L252R/G326A, AaaPT-NGR: C176N/S178G/L253R, AaaPT-NGRA: C176N/S178G/L253R/G327A, AmaPT-NGSRG: Y193S/S176N/S178G/M253R/M327G, AmaPT-NGSRA: Y193S/S176N/S178G/M253R/M327A.

Structural models of AfaPT-WT, AaaPT-WT, AmaPT-WT, and AmaPT-SLG, generated using AlphaFold3, were subjected to structural alignment. The active sites of AfaPT-WT and AaaPT-WT closely resemble that of AtaPT-WT, with key residues C175, S177, L252, and G326 well conserved (Figure 4b), suggesting that introduction of these residues via site-directed mutagenesis could confer similar catalytic properties. In contrast, comparison of AtaPT-WT with AmaPT-WT and AmaPT-SLG revealed that residues Y193, M253, and M326 in AmaPT-WT adopt extended conformations that restrict the active-site cavity, thereby hindering accommodation of the geranyl moiety and resulting in negligible activity (Figure 4c). Substitution of these residues with their AtaPT counterparts alleviates this steric constraint, giving rise to the active AmaPT-SLG variant. Accordingly, AmaPT-SLG was selected as the template for further engineering.

To evaluate the transferability of the functional role of C175, the variants AfaPT-C175A, AfaPT-C175H, AaaPT-C175A, AaaPT-C175H, AmaPT-ASLG, and AmaPT-HSLG were constructed and characterized. All variants exhibited enhanced catalytic performance (Figure 4d). AfaPT-C175A showed a 2.1-fold increase in activity with 86.7% selectivity toward **3a**, whereas AfaPT-C175H achieved a 4.5-fold increase with 93.5% selectivity toward **4a**. AaaPT-C175A displayed modest improvements (activity 23.5%, selectivity 56.7% toward **3a**), while AaaPT-C175H exhibited a 6.1-fold increase in activity with 94.1% selectivity toward **4a**. For AmaPT, the ASLG variant showed moderate enhancement in activity (20.3%) alongside a marked increase in selectivity (71.5%) toward **3a**, whereas AmaPT-HSLG exhibited a 3.1-fold increase in activity with a more modest improvement in selectivity toward **4a** (Figure 4d). Collectively, these results demonstrate that C175 acts as a transferable switch that simultaneously modulates both catalytic activity and selectivity across homologous PTs.

Building on these improvements, additional mutations—AfaPT-NGR, AfaPT-NGRA, AaaPT-NGR, AaaPT-NGRA, AmaPT-NGSRG and AmaPT-NGSRA—were introduced to further enhance catalytic performance toward **2a** (Figure 4e). The activities of AfaPT-NGR and AfaPT-NGRA increased by nearly 12-fold relative to AfaPT-WT, with selectivities improved to 79.0% and 93.3%, respectively. Similarly, AaaPT-NGR and AaaPT-NGRA showed 3.3-fold and 2.0-fold increases in activity, with selectivities of 84.9% and 80.8%, respectively. For AmaPT, NGSRG exhibited a 3.3-fold increase in activity with 65.9% selectivity, whereas NGSRA achieved higher selectivity (81.2%) at the expense of activity (2.2%).

Collectively, these results demonstrate that the mechanistic principles derived from AtaPT can be effectively transferred to homologous enzymes, indicating that the selectivity switch is broadly conserved rather than unique. The identification of key residues and engineered variants not only enables practical biocatalytic production but also establishes a general framework for the rational design and discovery of improved prenyltransferases.

### 2.6 Substrate Scope Exploration

Owing to the excellent substrate promiscuity of AtaPT [11] and the high catalytic performance of the engineered variants, we next evaluated their substrate scope. In addition to **1a**, 59 substrates were tested using WT, NGR, NGRA, C175A, and C175H as catalysts. Based on structural features, these substrates were classified into six groups: (I) resorcinol derivatives with alkyl or cyano substituents (**1b**–**1n**); (II) resorcinol derivatives bearing acyl groups (**1o**–**1ae**); (III) catechol derivatives with acyl substitution (**1af**–**1ah**); (IV) phloroglucinol- or pyrogallol-based compounds (**1ai**–**1aq**); (V) phenol derivatives with acyl substituents (**1ar**–**1bc**); and (VI) other small phenolic compounds (**1bd**–**1bh**) (Figure 5).

**FIGURE 5.**
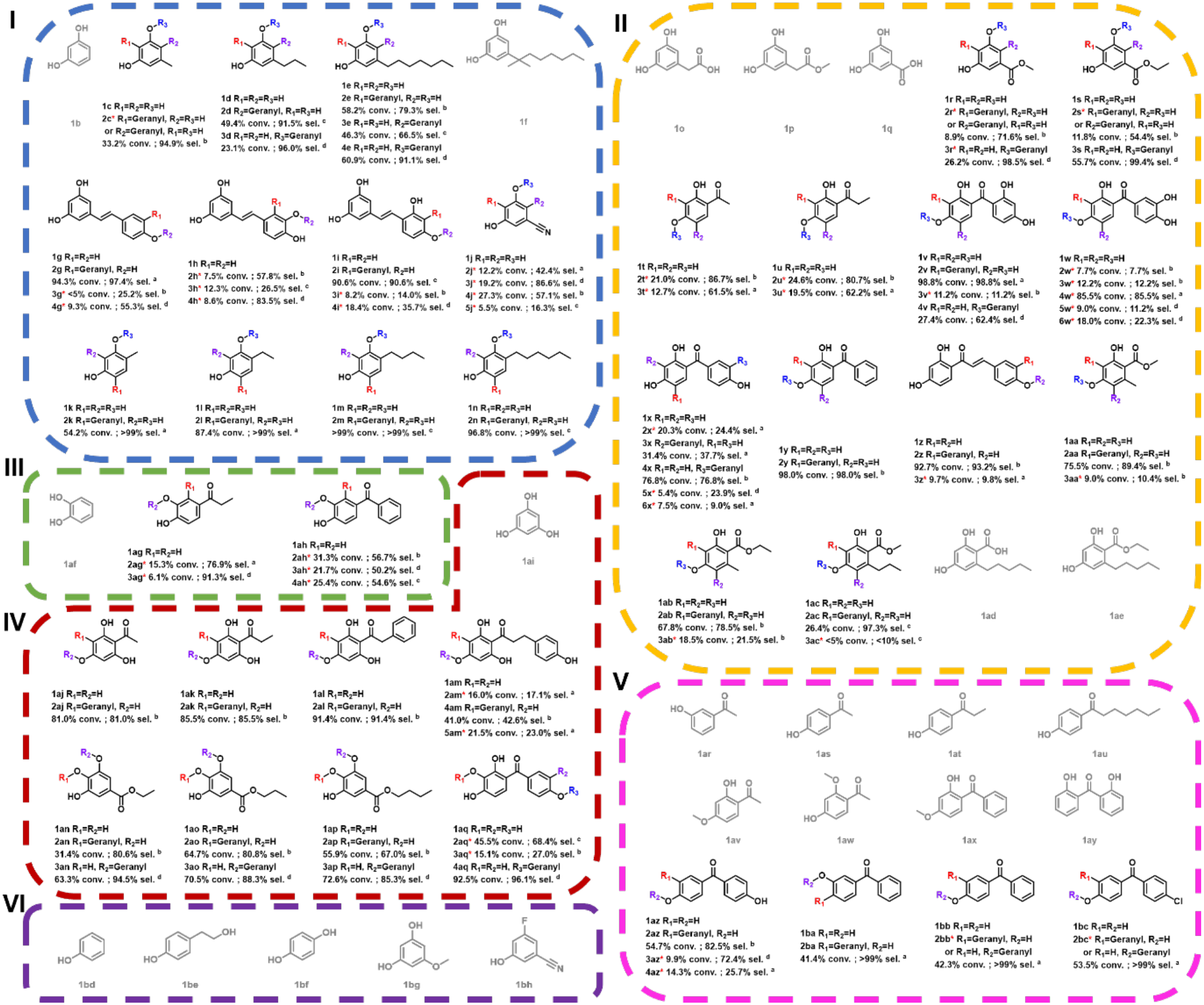
Substrate scope exploration. Structures of the tested substrates and corresponding geranylated products. Biotransformations were performed with 2 mM aromatic acceptor substrate, 4 mM prenyl donor GPP, 5 mM CaCl₂, and whole cells (OD₆₀₀ = 40-60) in a 300 µL reaction volume containing 2% (v/v) DMSO and 50 mM Tris-HCl buffer (pH 7.5). Reactions proceeded for 10-12 h, and products were isolated and characterized by HRMS and ^1^H and ^13^C NMR spectroscopy. The indicated conversion rate and selectivity reflect the response of the best variant, where WT (a), NGR (b), C175A (c), and C175H (d).

Overall, excluding NGRA—which nearly abolished both activity and selectivity toward all additional substrates—the other three variants exhibited substrate specificity similar to that of the WT, retaining high promiscuity by accepting 37 of the 59 tested substrates (Figures S19-S56). Inactive substrates were mainly found in groups V and VI. Structural analysis reveals two key determinants of substrate specificity. For group V, the acyl substituent must be larger than a phenyl group, and a free phenolic hydroxyl group cannot be positioned ortho to the acyl group, likely due to steric constraints and inhibition of phenolic deprotonation. For group VI, all substrates were inactive; comparison with other groups suggests that, in the absence of an acyl group, at least two phenolic hydroxyl groups are required for substrate recognition. Furthermore, the inactivity of **1b**, **1af**, **1ai**, and **1bf**, together with similar trends observed for **1f** (group I) and **1o**, **1p**, **1q**, **1ad**, and **1ae** (group II), indicates that, beyond the phenolic core, appropriately sized alkyl, acyl, or ester substituents are essential for catalytic activity. Specifically, the inactivity of **1f** is likely attributed to severe steric hindrance introduced by its highly branched *α*-dimethyl group, which prevents proper accommodation within the active site. The loss of activity for **1o** and **1p** indicates that their truncated side chains fail to provide the essential hydrophobic interactions required for substrate anchoring. Similarly, the inactivity of **1ad** and **1ae** suggests poor active-site compatibility, likely due to electrostatic repulsion from the free carboxyl group (**1ad**) and spatial clashes from the bulky ethyl ester (**1ae**). Collectively, these results demonstrate that precise spatial and polar matching of the substituents is critical for effective catalytic turnover.

Among the 37 active substrates, WT exhibited the highest activity and selectivity for 15 substrates (conversion > 55 %, selectivity > 85 %), preferentially catalyzing geranylation at the *C*3′ position or *C*6 position (**1g, 1i, 1k**-**1n** in Class I; **1v**, **1w**, **1y**, **1z**–**1ab** in Class II; and **1ba**-**1bc** in Class V). It showed moderate activity with good selectivity for 3 substrates (conversion > 16 %, selectivity > 89 %), favoring geranylation at the *C*4 position or *C*6 position (**1c**, **1d** in Class I and **1ac** in Class II), and moderate activity but poor selectivity for the remaining 19 substrates (conversion > 10 %), including **1e**, **1h**, and **1j** (Class I); **1r**–**1u** and **1x** (Class II); **1ag** and **1ah** (Class III); **1aj**–**1aq** (Class IV); and **1az** (Class V) (Figures 5 and S57). The NGR and C175A variants displayed complementary behavior relative to WT, showing enhanced activity while retaining selectivity toward substrates such as **1b**, **1c**, **1i**, **1m**, **1n**, **1y**, and **1z**-**1ac** compared to the WT. Notably, NGR significantly improved both activity and selectivity for multiple products (**2e**, **4x**, **2y**, **2aj**, **2ak**, **2al**, **4am**, **2an**, **2ao**, **2ap**, **and 2az)** to 2-*C* geranylation (conversion > 41 %, selectivity > 43 %), consistent with the model reaction. The exceptions were substrate **1an**-**1ap** and **1ba**, where the hydroxyl group occupies the *C*2 position, forcing geranylation to occur at the hydroxyl (**2an**-**2ap**) or a neighboring site (**2ba**) (Figures 5 and S58). C175A promoted efficient production of **3a**, **2d**, **3e**, **2m**, and **2n** (conversion > 46 %, selectivity > 66 %). Among these, **3a**, **2d**, and **3e** were substituted at the *C*4 position, whereas **2m**-**2n**, similar to **2k**-**2l**, were substituted at the *C*6 position due to changes in the orientation of the carbon chain (Figures 5 and S58). Notably, NGR and C175A mainly enhanced the formation of the WT’s major products, while C175H was characterized as the unique *O*-geranyltransferase, exhibiting rather different selectivity compared to the other variants. C175H tended to generate *O*-geranylated products; for example, **4a**, **3d**, **4e**, **3s**, **3an**, **3ao**, and **3ap** were substituted at the C3-OH position (conversion > 23 %, selectivity > 85 %). Likely due to neighboring group effects and the influence of B-ring hydroxyls, substitution of **4v** occurred at the 1-OH position, whereas **4aq** was substituted at the 4’-OH of the B-ring (Figures 5 and S58). Structurally, these cases are analogous, as **1v** has a symmetrical structure. However, for substrates **1c**, **1k**-**1n**, **1y**-**1ac**, **1ba**-**1bc**, and the *O*-geranylation activity of C175H, no noticeable activity was observed. Moreover, compared with the WT, the overall catalytic activity was significantly decreased.

Overall, for substrates in the substrate scope, the complementary activities and selectivities of the AtaPT-WT and the mutants enable the corresponding formation of multiple products using the appropriate catalysts. This also suggests that the AtaPT variants can be combined to construct a platform for the production of geranylation products.

### 2.7 Coupling the isopentenol utilization pathway with prenyltransferase for prenylflavonoid biosynthesis

The isopentenol utilization pathway (IUP) converts exogenously supplied isoprenol or prenol into dimethylallyl diphosphate (DMAPP) or isopentenyl pyrophosphate (IPP) via sequential phosphorylation, providing a streamlined route to isoprenoid precursors. Coupling the IUP with PTs has been widely employed for the efficient production of diverse prenylated compounds [41-44]. In our previous work, we established a whole-cell system that generates GPP from geraniol via the IUP to supply downstream PTs for the biosynthesis of geranylated products, although the overall efficiency remained relatively low [35].

To further improve geranylation efficiency, we investigated the GPP biosynthetic pathway in greater detail. In principle, the conversion of geraniol to GPP proceeds through two sequential phosphorylation steps: geraniol is first phosphorylated to geranyl phosphate (GP), which is subsequently converted to GPP. Given the excellent catalytic performance of AtaPT-M7 toward naringenin, we selected AtaPT-M7 and naringenin as a model system to evaluate the efficiency of GPP regeneration [35]. THAIPK mutants have been reported to catalyze the second phosphorylation step [45]. Among them, comparison of THAIPKG73 (THAIPK-Y70A/V73A/I140A) and THAIPKG130 (THAIPK-Y70A/V130A/I140A) revealed that THAIPKG73 exhibits superior catalytic activity (Figure S59).

After identifying the optimal isopentenyl phosphate kinase, we next investigated the enzyme responsible for the first phosphorylation step of geraniol. Diacylglycerol kinase A (DGKA) from *Streptococcus mutans* has recently been reported to function as a decaprenol kinase capable of phosphorylating geraniol to geranyl phosphate (GP) [46, 47], providing a useful template for mining potential candidates. Accordingly, homologous enzymes, including EcDGKA from *E. coli* and BsDGKA from *Bacillus subtilis*, were selected. In addition, the engineered variant SmUdpKM2 (SmUdpK-S47A/L124A), previously reported to exhibit enhanced activity toward prenol substrates [48], was also evaluated. A series of plasmid constructs—pCDF-THAIPKG73-SmUdpK, pCDF-THAIPKG73-SmUdpKM2, pCDF-THAIPKG73-EcDGKA, and pCDF-THAIPKG73-BsDGKA—were generated and assessed, among which pCDF-THAIPKG73-SmUdpK showed the highest catalytic efficiency (Figure S59).

Based on these results, pCDF-THAIPKG73-SmUdpK and pET28a-M7 were selected for further optimization. Key parameters, including expression temperature, inducer concentration, reaction temperature and pH, catalyst ratio, and the concentrations of enzyme, geraniol, Mg^2+^, and DMSO, were systematically examined. The optimal conditions were identified as follows: THAIPKG73-SmUdpK : M7 = 1:4, OD_600_ = 100, 50 mM Tris–HCl buffer (pH 7.5), 2 mM acceptor, 4 mM geraniol, and 4% (v/v) DMSO, with incubation at 25 °C (Figures S60-S61). Notably, geraniol concentration had a pronounced effect on catalytic performance, with excessive substrate levels leading to severe inhibition (Figure S62). Under these optimized conditions, the reaction reached a plateau after 12 h, and the efficiency of M7-catalyzed geranylation of naringenin exceeded 90%, comparable to that achieved by direct GPP supplementation, indicating efficient operation of the GPP regeneration pathway.

The versatility of the cascade platform was further validated across multiple mutants. A set of 10 substrates was evaluated in a 50 mL cascade system, with WT applied for the synthesis of **2g** and **2z**, NGR for **2y**, **2aa**, and **2ab**, C175A for **2i**, **2m**, and **2v**, and C175H for **3ap** and **4aq**. All corresponding products were obtained in satisfactory yields: **2g** (47.3 %), **2z** (48.5 %), **2y** (63.4 %), **2aa** (40.5 %), **2ab** (44.3 %), **2i** (49.9 %), **2m** (38.5 %), **2v** (66.9 %), **3ap** (48.6 %), and **4aq** (42.7 %). These results demonstrate the robustness and broad applicability of the cascade system (Figure 6).

**FIGURE 6.**
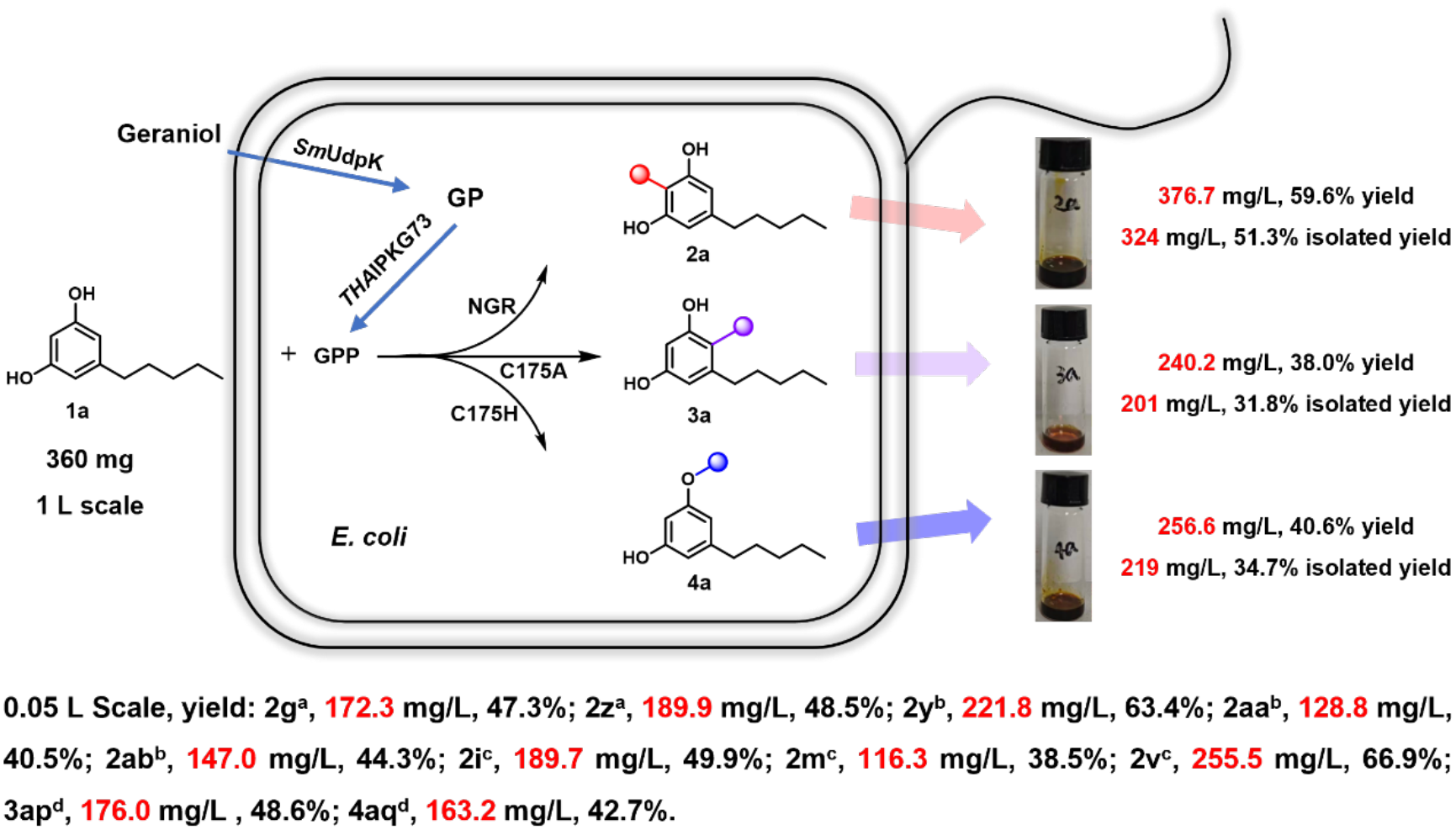
Biocatalytic cascade for the synthesis of geranylated olivetol derivatives. *E. coli* cells co-expressing the donor supply system and AtaPT mutants (1:4 ratio) were resuspended in 50 mM Tris–HCl buffer (pH 7.5) and adjusted to an OD₆₀₀ of 100. After complete lysis, the reactions were carried out using the resulting lysate. Substrates were added to final concentrations of ∼2 mM olivetol and 4 mM geraniol in a total volume of 1 L, and bioconversions were performed at 25 °C with shaking at 220 rpm for 12 h. Symbols denote the AtaPT variants used: ^a^ WT; ^b^ NGR; ^c^ C175A; and ^d^ C175H. THAIPKG73 refers to THAIPK-Y70A/V73A/I140A.

Furthermore, preparative-scale reactions of **1a** were conducted in 1 L systems, affording isolated product titers of 324 mg/L **2a** (NGR), 201 mg/L **3a** (C175A), and 219 mg/L **4a** (C175H), respectively (Figure 6). Collectively, these results highlight the scalability and generality of the engineered geranylation platform, providing a robust and flexible strategy for the biosynthesis of structurally diverse geranylated compounds.

## 3 Conclusion

Understanding the selectivity of aPTs is critical for reconciling their broad substrate scope with poor regio- and chemoselectivity, yet this remains a major challenge. In this study, we elucidated the mechanistic basis of regio- and chemoselective geranylation of resorcinol catalyzed by AtaPT through directed evolution, identifying a unique single-site switch (C175). Engineering this site yielded a series of variants (C175A, C175H, NGR, and NGRA) that collectively enable access to all possible mono-geranylation products. Structural characterization and computational simulations further revealed the regulatory mechanism of this residue in controlling selectivity and activity: C175 functions as a crucial spatial and steric gatekeeper. The size, polarity, and steric profile of the residue at this position precisely dictate the binding depth and orientation of the phenolic acceptor relative to the prenyl donor. Specifically, smaller side chains (C175A) accommodate lateral substrate binding to favor *C*4 geranylation; bulky residues (C175H) induce substrate rotation via steric hindrance to drive *O*-geranylation; and targeted polar substitutions (C175N) facilitate deeper pocket penetration for selective *C*2 geranylation. Importantly, the generality of this mechanism was validated by rational engineering of three additional aPTs, enabling tunable selectivity and improved catalytic performance.

Leveraging the broad substrate scope and enhanced selectivity of the engineered variants, 37 structurally diverse polyphenols were efficiently geranylated. By integrating an *in situ* GPP regeneration system, 10 geranylated products were synthesized at semi-preparative scale (50 mL), while preparative-scale reactions (1 L) afforded three geranylated olivetol derivatives with titers exceeding 200 mg/L. These results demonstrate that the engineered cascade system provides a robust and scalable platform for the biosynthesis of geranylated polyphenols.

Overall, this work establishes an efficient and economical single-site strategy for controlling chemo- and regioselectivity as well as catalytic activity in aPTs, and elucidates the underlying molecular mechanism. These insights offer a foundation for the rational design and customization of aPTs for selective geranylation.

## 4 Methods and Materials

### 4.1 Reagents and instruments

All chemical reagents were purchased from Aladdin, Macklin Biochemical, Meryer (Shanghai) Chemical Technology Co., 9ding Chemistry, Bide Pharm Tech, Shanghai Yuan Ye Biotechnology Co., or Sangon Biotech (Shanghai, China) unless otherwise stated. The KOD One TM PCR master Mix polymerase was obtained from Toyobo Ideas & Chemistry. DpnI was purchased from Abclonal Technology. The ClonExpress MultiS One-Step Cloning Kit was obtained from Vazyme Biotech. Primer synthesis and DNA sequencing were conducted at Sangon Biotech Company (Shanghai, China). Micro-spectrophotometer (Allsheng, Nano-300), ultrasonic homogenizer with noise isolating chamber (SCIENTZ-IID), freeze drier (SCIENTZ-10ND), centrifuge (Thermo Fisher, (Sorvall ST 16R) and PICO 17), PCR Amplifier (BIO-RAD, T100TM Thermal Cycler), mini shaker (Allsheng, MSC-100), shaker (Minquan, MQD-BIR). Shimadzu Nexis LC 20AT equipped with an electrospray ionization (ESI) source was used for the measurement of substrates and products. High resolution mass spectrometry (HRMS) detection was performed by X500R QTOF from AB SCIEX. The prenylated products were isolated and purified by semi-preparative HPLC on an LC-8A instrument (Shimadzu). Products were characterized by 1HNMR and 13C NMR in DMSO-d6 on AV2 500MHz Bruker spectrometer. Chemical shifts (δ) were given in ppm and coupling constants (J) were given in hertz (Hz).

### 4.2 Heterogonous expression and purification of PTs

The AtaPT gene was synthesized by Genewiz (China) and cloned into the *Nde*I/*Xho*I sites of the pET28a vector. The resulting recombinant plasmid (pET28a-AtaPT) was transformed into *E. coli* BL21 (DE3) for heterologous expression.

A seed culture (5 mL) was grown overnight in Luria–Bertani (LB) medium supplemented with 50 μg/mL kanamycin at 37 °C with shaking (220 rpm). Subsequently, 1% (v/v) of the seed culture was inoculated into 500 mL TB medium containing 50 μg/mL kanamycin and cultivated at 37 °C and 220 rpm in 2 L shake flasks. When the culture reached an OD₆₀₀ of 0.8–1.0, protein expression was induced with 0.2 mM IPTG, followed by incubation at 18 °C and 220 rpm for 16 h.

Cells were harvested by centrifugation (6,438 × g, 3 min, 4 °C) and resuspended in binding buffer (50 mM Tris–HCl, 5 mM CaCl₂, pH 7.5). Cells were disrupted by sonication in an ice bath, and cell debris was removed by centrifugation (15,480 × g, 4 °C, 1 h). The supernatant was filtered through a 0.45 µm membrane and loaded onto an ÄKTA avant 25 system equipped with a 5 mL His Trap™ HP column. Proteins were purified using a linear imidazole gradient (10-500 mM) in 50 mM Tris-Cl buffer (pH 7.5) containing 5 mM CaCl₂, following the manufacturer’s instructions.

### 4.3 Construction and screening of AtaPT mutant libraries

All strains, plasmids, and primers used in this study are listed in Tables S2 and S3. PCR amplification (Table S4) was performed using KOD One™ PCR Master Mix. After sequence verification, recombinant plasmids were transformed into *E. coli* BL21 (DE3) for heterologous protein expression as described above.

Single-site saturation mutagenesis (SSM) and iterative saturation mutagenesis (ISM) were carried out using primers containing NNK codons at the target positions. To ensure sufficient library coverage, 92 colonies were screened for each mutagenized site. Colonies were cultivated in 96-deep-well plates containing 0.3 mL LB medium supplemented with 50 μg/mL kanamycin at 37 °C overnight. Subsequently, 0.7 mL TB medium containing 50 μg/mL kanamycin and 0.2 mM IPTG was added, and cultures were incubated at 18 °C for 16 h.

Cells were harvested and resuspended in 0.2 mL reaction buffer (2 mM substrate **1a**, 2 mM GPP, 50 mM Tris–HCl, 5 mM CaCl₂, pH 7.5) and incubated at 37 °C for 24 h with shaking. Reactions were quenched by the addition of 0.6 mL methanol.

Relative activities were evaluated by whole-cell biotransformations in a total volume of 2 mL reaction buffer (2 mM **1a**, 2 mM GPP, 50 mM Tris–HCl, 5 mM CaCl₂, pH 7.5) at 37 °C for 24 h, using the wild-type enzyme as a control. After centrifugation (12,000 × g, 15 min), supernatants were analyzed by HPLC–UV using a Shimadzu Shim-pack GIS C18 column (4.6 × 250 mm, 5 μm) at a flow rate of 1 mL/min and a column temperature of 30 °C. Gradient elution programs are provided in Table S5.

### 4.4 Molecular docking of AtaPT-WT and substrate

Molecular docking was performed using AutoDock Vina. The crystal structure of wild-type AtaPT (PDB ID: 5KCG) was first imported and prepared as a PDBQT file. The GPP molecule was similarly processed: the initial MOL file was converted to a PDBQT format after energy minimization. Docking of GPP into the protein structure was carried out using Vina, with the search space defined to encompass the prenyl donor binding pocket, including residues K95, R104, K188, R264, K266, N328, Q342, and N411. The docking pose with the lowest binding energy was selected and further analyzed in PyMOL. Subsequently, olivetol (**1a**) was docked into the protein using Vina, targeting the aromatic substrate-binding pocket defined by residues I194, L204, L249, L252, A270, F272, L323, and W397. Docking results were filtered in PyMOL by identifying the pose with the shortest C-C distance between the reactive centers of GPP and **1a**, indicating potential for productive prenylation. Finally, residues within a 5 Å radius of each ligand were selected and visualized to highlight the local binding environment (Figure S4).

### 4.5 Determination of kinetic parameters

Kinetic analyses were performed for AtaPT-WT, AtaPT-NGR, AtaPT-NGRA, AtaPT-C175A, and AtaPT-C175H using GPP as the prenyl donor and **1a** as the acceptor substrate. Enzymatic assays were conducted in 100 µL reaction volumes containing 50 mM Tris-HCl buffer (pH 7.5), 5 mM CaCl₂, 10 µM enzyme, 2 or 4 mM GPP (saturating), and varying concentrations of **1a** (0.5–4.0 mM).

Reactions were incubated at 37 °C for 10–20 min and quenched by the addition of 300 µL methanol, followed by centrifugation (13,800 × g, 15 min). The resulting supernatants were analyzed by HPLC as described above.

All measurements were performed in triplicate. Michaelis–Menten parameters (*K*ₘ) were obtained by nonlinear regression fitting, and turnover numbers (*k*_cat_) were calculated accordingly (Table S8 and Figure S14).

### 4.6 Structure determination of AtaPT variants

The AtaPT variants (NGR, NGRA, C175A and C175H (11-424)), each bearing an *N*-terminal 6×His tag, were overexpressed in *E. coli* BL21 (DE3) cells. Protein expression was induced with 0.4 mM IPTG when cultures reached an OD_600_ of 0.8, followed by incubation at 18 °C for 12 h. Cells were harvested and lysed in lysis buffer containing 25 mM Tris-HCl (pH 8.0), 1 M NaCl, 25 mM imidazole, supplemented with 0.5 mM *β*-mercaptoethanol (*β*-ME), and 1 mM phenylmethylsulfonyl fluoride (PMSF). The His-tagged fusion proteins were purified using Ni²⁺-affinity chromatography. The column was washed with lysis buffer, and proteins were eluted with elution buffer containing 25 mM Tris-HCl (pH 8.0), 1 M NaCl, 250 mM imidazole. Further purification was performed by size-exclusion chromatography using a Superdex 200 column (GE Healthcare) pre-equilibrated with buffer containing 25 mM Bis-Tris (pH 6.5), 200 mM NaCl, 5% (v/v) glycerol, and 3 mM dithiothreitol (DTT). The purified proteins were concentrated to 25 mg/mL and stored at -80 °C.

The apo AtaPT variants crystals were grown in the crystallization conditions of 0.1 M Bis-Tris (pH 6.5), 0.2 M ammonium sulfate, and 15-20% (w/v) PEG 3350 using hanging-drop vapour diffusion at room temperature. To obtain the crystals of ternary complex, the apo crystals were first soaked with prenyl donor compounds (GSPP or GPP) at a final concentration of 5 mM, and then soaked with compound **1a** at a final concentration of 25 mM. Prior to diffraction experiments, all of the crystals were soaked in the reservoir buffer supplemented with 25% (v/v) glycerol. X-ray diffraction data were collected at the BL19U1 beamline of the Shanghai Synchrotron Radiation Facility (SSRF). The datasets were indexed, integrated and scaled using HKL3000 package. The structures were solved by molecular replacement using PHASER with the structure of AtaPT-WT (PDB: 5KCG) as the searching model. Following iterative manual rebuilding and refinement were performed with the use of COOT and PHENIX, respectively.

### 4.7 Reaction condition optimization

All optimization experiments were performed in triplicate on an analytical scale (300 µL) using whole-cell catalysts, with GPP as the prenyl donor and **1a** as the acceptor substrate. Unless otherwise stated, reactions were conducted in 50 mM Tris–HCl buffer (pH 7.5) containing 5 mM CaCl₂, based on previously reported protocols.

To determine the optimal reaction temperature, assays were carried out at 25, 30, 37, 45, and 50 °C for 24 h. The effect of pH was evaluated in Tris-HCl buffers ranging from pH 7.0 to 9.0 under the optimal temperature conditions. The influence of the co-solvent was assessed by varying DMSO concentrations (2-10%, v/v) under otherwise identical conditions.

The effect of cell density was investigated by performing reactions at OD₆₀₀ values of 20, 40, 60, 80, and 100. Product formation (**2a**, **3a**, and **4a**) increased with cell density up to OD_600_ = 40-60 and decreased at higher cell densities (OD₆₀₀ = 100).

Time-course experiments were conducted under optimized conditions (30-45 °C, 50 mM Tris-HCl, pH 7.5, 2% DMSO), with samples taken at 2, 4, 6, 8, 10, 12, and 24 h. Reactions were quenched by addition of 0.9 mL methanol, followed by centrifugation, and the supernatants were analyzed by analytical HPLC as described previously (Table S6 and Figures S6–S10).

### 4.8 Preparation of E. coli whole cell catalyst harboring AtaPT variants

Following protein expression, cells were harvested by centrifugation (6,438 × g, 2 min, 4 °C) and resuspended in binding buffer (50 mM Tris–HCl, 5 mM CaCl₂, pH 7.5) to an OD_600_ of 40–60. The resulting cell suspensions were aliquoted, and the wet cell pellets were stored at −80 °C until further use.

### 4.9 Construction and characterization of other PTs variants

AtaPT was used as a query for BLAST searches against the NCBI database to identify homologous prenyltransferases. Three enzymes with varying sequence identities were selected: AfaPT (94.81%, GenBank: KAL5359867.1), AaaPT (86.7%, GenBank: XP_073546247.1), and AmaPT (56.87%, GenBank: XP_045942482.1). The corresponding genes were synthesized (General Biosystems, Anhui, China) and cloned into the *Nde*I/*Xho*I sites of the pET28a vector. The resulting plasmids (pET28a-AfaPT, -AaaPT, and -AmaPT) were transformed into *E. coli* BL21(DE3) for heterologous expression.

For protein expression, *E. coli* cultures were grown overnight in 5 mL LB medium supplemented with kanamycin (50 μg/mL) at 37 °C with shaking (220 rpm). The cultures were then inoculated (1%, v/v) into 500 mL TB medium containing kanamycin (50 μg/mL) and cultivated at 37 °C (220 rpm) in 2 L flasks. When OD_600_ reached 0.8-1.0, protein expression was induced with 0.2 mM IPTG, followed by incubation at 18 °C for 16 h.

Whole-cell biotransformation assays were carried out in 300 µL reaction mixtures containing 50 mM Tris-HCl buffer (pH 7.5), 5 mM CaCl₂, 4 mM geranyl diphosphate (GPP), 2 mM acceptor substrate (**1a**), 2% (v/v) DMSO, and enzyme preparations adjusted to OD_600_ = 40-60. Reactions were incubated at 30-45 °C for 10-12 h and quenched by the addition of 900 µL methanol. After centrifugation (13,800 × g, 15 min), the supernatants were analyzed by HPLC.

### 4.10 HPLC-ESI and HRMS-based prenyltransferase activity assay

*In vitro* activity assays were performed in 300 µL reaction mixtures containing 4 mM GPP, 2 mM acceptor substrate, 2% (v/v) DMSO, and whole cells (OD_600_ = 40–60) in 50 mM Tris–HCl buffer (pH 7.5) supplemented with 5 mM CaCl₂. Reactions were incubated at 30-45 °C for 10-12 h and quenched by the addition of 900 µL methanol.

Following centrifugation (13,800 × g, 15 min), the supernatants were subjected to HPLC–ESI analysis and high-resolution mass spectrometry (HRMS). All assays were conducted in triplicate. Gradient programs used for analysis are provided in Table S5 and Figures S19–S56.

### 4.11 Product identification by NMR

Recombinant *E. coli* strains harboring pET28a-AtaPT-WT, pET28a-AtaPT-NGR, pET28a-AtaPT-C175A or pET28a-AtaPT-C175H were prepared and induced as described above. After induction, the whole cells were harvested by centrifugation and washed with 50 mM Tris-HCl buffer (pH 7.5) containing 5 mM CaCl₂. The cells were then resuspended in the same buffer, and the optical density was adjusted to OD_600_ = 40/60. The reaction mixture (total volume: 50 mL in a 250 mL Erlenmeyer flask) was prepared by adding 2 mM acceptor substrate and 4 mM GPP. Bioconversion was carried out at 30-45 °C with shaking for 10-12 h. After the reaction, the mixture was extracted with an equal volume of ethyl acetate and centrifuged at 6438 g for 10 min. The supernatant was collected and transferred to a separating funnel, where the aqueous phase was discarded, and the organic phase was retained. The solvent was removed under reduced pressure, and the resulting residue was dissolved in methanol and purified by reverse-phase semi-preparative HPLC. The purified products were dissolved in DMSO-*d₆* and characterized by ^1^H and ^13^C NMR spectroscopy.

### 4.12 Coupling of IUP with AtaPT-WT/NGR/C175A/C175H for the preparation of geranylated polyphenols using a lysate-based catalyst

The cascade biosynthesis of geranylated polyphenols was optimized using mixed whole-cell lysates of *E. coli* BL21(DE3) harboring pET28a-M7 and PCDF-SmUDPK-THAIPKG73, with naringenin as the model substrate. Key parameters were systematically evaluated, including expression temperature (18, 25, 30, or 37 °C), IPTG concentration (0.2, 0.5, 0.8, or 1 mM), reaction temperature and pH (20–45 °C; pH 7.0–9.0), enzyme lysate ratio (SmUDPK : THAIPKG73 = 1:5, 1:4, 1:2, 1:1, or 2:1), cell density (OD₆₀₀ = 20–150), Mg²⁺ concentration (0–10 mM), geraniol concentration (2–30 mM), and DMSO content (1–10% v/v). The reaction reached a steady state after approximately 12 h. Reactions were quenched by adding three volumes of methanol, followed by centrifugation at 13,800 × g for 15 min. Products were analyzed by HPLC, and data are reported as mean ± SD from three biological replicates (Figures S59-S62).

For strain cultivation, *E. coli* harboring the relevant plasmids was grown in TB medium supplemented with kanamycin (50 µg/mL) or spectinomycin (100 µg/mL) at 37 °C to an OD₆₀₀ of 0.8–1.0. Protein expression was induced with 0.2 mM IPTG, followed by incubation at 18 °C for 16 h. Cells were harvested by centrifugation (6,438 × g, 2 min, 4 °C) and resuspended in 50 mM Tris–HCl buffer (pH 7.5) to an OD₆₀₀ of 100.

Preparative-scale cascade reactions were conducted with 2 mM substrate (e.g., **1a**), 4 mM geraniol, and 4% (v/v) DMSO in a total volume of 1 L at 25 °C for 12 h. After completion, the reaction mixtures were extracted with two volumes of ethyl acetate and centrifuged (6,438 × g, 5 min). The organic layers were collected, concentrated under reduced pressure, and the residues were dissolved in dichloromethane (DCM) and purified by silica gel chromatography. The eluates were concentrated to afford the final products, which were further dried under reduced pressure.

Substrate scope evaluation was performed in 50 mL reactions using WT, NGR, C175A, or C175H in combination with SmUDPK-THAIPKG73. The WT system showed higher activity toward **2g** and **2z**; the NGR variant was used for the synthesis of **2y**, **2aa**, and **2ab**; C175A was employed for **2i**, **2m**, and **2v**; and C175H was used for the production of **3ap** and **4aq**.

## Acknowledgements

We acknowledge the financial support from the National Key Research and Development Program of China (No. 2023YFA0914100/2023YFA0914102), Key project at central government level: The ability establishment of sustainable use for valuable Chinese medicine resources (No. 2060302/2060302-2504-01), the National Natural Science Foundation of China (No. 22477110 and 22034002), the Fundamental Research Funds for the Central Universities (No. 226202400061 and 226202300100), Guangdong Major Project of Basic and Applied Basic Research (No. 2023B0303000026), the Noncommunicable Chronic Diseases-National Science and Technology Major Project (No. 2023ZD0505900), the Open Fund of Guangxi Key Laboratory of Tea Plant Germplasm Innovation and Resource Utilization (No. 2025ZD06) and Yuelushan Laboratory Talent Program (No. 2025RC2054). We thank the staffs from BL19U1 beamline and the National Facility for Protein Science in Shanghai (NFPS) and BL02U1 at Shanghai Synchrotron Radiation Facility, for assistance during data collection. This article is dedicated to the memory of Prof. Manfred T. Reetz with deep sorrow.

